# Estimation of genetic correlation using linkage disequilibrium score regression and genomic restricted maximum likelihood

**DOI:** 10.1101/194019

**Authors:** Guiyan Ni, Gerhard Moser, Schizophrenia Working Group of the Psychiatric Genomics Consortium, Naomi R. Wray, S. Hong Lee

## Abstract

Genetic correlation is a key population parameter that describes the shared genetic architecture of complex traits and diseases. It can be estimated by current state-of-art methods, i.e. linkage disequilibrium score regression (LDSC) and genomic restricted maximum likelihood (GREML). The massively reduced computing burden of LDSC compared to GREML makes it an attractive tool, although the accuracy (i.e., magnitude of standard errors) of LDSC estimates has not been thoroughly studied. In simulation, we show that the accuracy of GREML is generally higher than that of LDSC. When there is genetic heterogeneity between the actual sample and reference data from which LD scores are estimated, the accuracy of LDSC decreases further. In real data analyses estimating the genetic correlation between schizophrenia (SCZ) and body mass index, we show that GREML estimates based on ~150,000 individuals give a higher accuracy than LDSC estimates based on ~400,000 individuals (from combined meta-data). A GREML genomic partitioning analysis reveals that the genetic correlation between SCZ and height is significantly negative for regulatory regions, which whole genome or LDSC approach has less power to detect. We conclude that LDSC estimates should be carefully interpreted as there can be uncertainty about homogeneity among combined meta-data sets. We suggest that any interesting findings from massive LDSC analysis for a large number of complex traits should be followed up, where possible, with more detailed analyses with GREML methods, even if sample sizes are lesser.

## MAIN TEXT

Genetic correlation is a key population parameter that describes the shared genetic architecture of complex traits and diseases ^1-3^. The genetic correlation is the additive genetic covariance between two traits scaled by the square root of the product of the genetic variance for each trait (i.e., the geometric mean of the trait variances). The sign of the correlation shows the direction of sharing, and the parameter definition is based on genetic variants across the allelic spectrum. Methods to estimate genetic correlation based on genetic covariance structure are well established for both quantitative and disease traits, e.g. (restricted) maximum likelihood for linear mixed models (LMM) ^4-6^. Genetic covariance structure can be derived from phenotypic records using pedigree information in twin or family-based designs ^7^. Recently, genome-wide single nucleotide polymorphism (SNP) data have been used to construct a genomic relationship matrix for the genetic covariance structure in LMM that captures the contribution of causal variants that are in linkage disequilibrium (LD) with the genotyped SNPs^4; 8; 9^. Such estimates assume that the genetic correlation estimated from common SNPs is representative of the parameter that depends on all genetic variants; this seems like a reasonable assumption.

In contrast to the genomic restricted maximum likelihood (GREML) approach, a linkage disequilibrium score regression (LDSC) ^10; 11^ method does not require individual-level genotype data but instead uses GWAS summary statistics, regressing association test statistics of SNPs on their LD scores. The LD score of a SNP is the sum of LD *r*^2^ measured with all other SNPs, and can be calculated in a reference sample of the same ethnicity when individual genotype data are not available for the GWAS sample, under the assumption that the GWAS sample has been drawn from the same ethnic population as the reference sample used to calculate the LD scores. The method exploits the relationship between association test statistic and LD score expected under polygenicity. Because of this simplicity, and the massively reduced computing burden in terms of memory and time, it is feasible for LDSC to be applied to a large number of multiple traits, e.g. Bulik-Sullivan et al. ^11^, Zheng et al. ^12^, Finucane et al. ^13^.

Given the attractiveness of LDSC for a massive analysis of many sets of GWAS summary statistics, it has been widely used in the community. However, genetic correlations estimated by LDSC are often reported without caution although the approach is known to be less accurate, compared to GREML^11^. In fact, the accuracies of LDSC estimates have not been thoroughly studied.

In this report, we compare both the bias (difference between the simulated true value and estimated value) and accuracy (i.e. magnitude of the standard error of an estimate, SE) between GREML and LDSC for estimation of genetic correlation. We find that both methods show little evidence of bias. However, LDSC is less accurate as reported in Bulik Sullivan et al.^11^, with SE at least more than 1.5-fold higher than that of GREML regardless of the number of samples in data used to estimate the genetic correlation. When decreasing the number of SNPs, the accuracy of LDSC decreases further. When increasing the degree of genetic heterogeneity between the actual sample and reference data from which LD scores are estimated, the SE of LDSC estimates are up to 3-fold larger than those of the GREML estimates. We also show that GREML is more accurate in genomic partitioning analyses over LDSC or stratified LDSC (sLDSC). In genomic partitioning analyses the genetic parameters are estimated for genomic subsets defined by user-specified annotations. In analyses of real data, we show that GREML is more accurate and powerful, e.g. GREML estimates based on ~ 150,000 individuals give a higher accuracy than LDSC estimates based on 400,000 individuals in estimating genetic correlation between schizophrenia (SCZ) and body mass index (BMI) (-0.136 (SE=0.017) and p-value=4.54E-15 for GREML vs. -0.087 (SE=0.019) and p-value=4.91E-06 for LDSC). In these analyses, the GREML estimate is based on UK sample only whereas the LDSC estimate is based on combined meta-data sets among which there is uncertainty about homogeneity. Furthermore, a GREML genomic partitioning analysis reveals that the genetic correlation between SCZ and height is significantly negative for regulatory regions, which is less obvious by LDSC both when using whole-genome or partitioned estimates of genetic correlation.

In the main methods, we used GREML^14; 15^ and LDSC^10; 11^ to compare their estimates of genetic correlation using simulated as well as real data. Simulations were based on UK Biobank imputed genotype data (UKBB^16^) after stringent quality control (QC) (see Supplemental Methods). We calculated a ratio of empirical SE and its 95% confidence interval (CI) to assess the accuracy of the methods for each set of simulated data. The 95% CIs of SE were estimated based on the delta method^17^. When estimating genetic correlation using simulated phenotypes based on UKBB genotype data we found that the estimates were unbiased for both GREML and LDSC (Figure S1), but the SE of GREML was at least 1.5 times smaller than that of LDSC (Figure 1). The ratio of the empirical SE from LDSC to GREML was increased up to 3.5-fold when using a smaller number of SNPs (Figure 1). All values of the ratio were significantly different from 1. It is notable that the SE of GREML estimates showed almost no difference across different numbers of SNPs whereas that of LDSC estimates gradually increased with a smaller number of SNPs (Figure S2). The ratio was invariant to sample size (Figure S3). As expected, when using the intercept constrained to zero, LDSC estimates were substantially biased when there were overlapping samples (Figure S4). We also explored alternative genetic architectures (Figure S5), which consistently showed that GREML gives a smaller SE than LDSC in any scenario.

**Figure 1.**
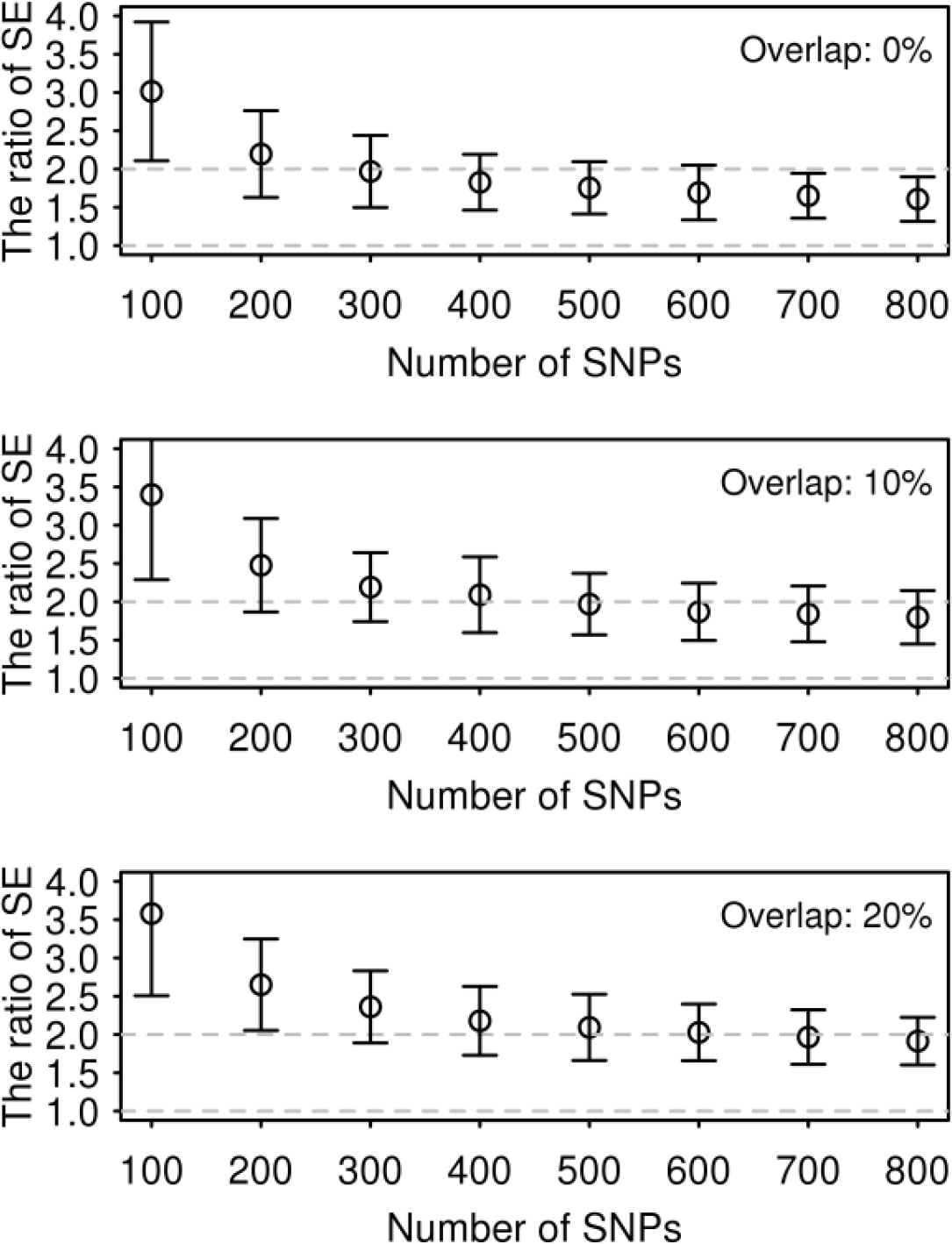
The ratio of SE of LDSC estimate to that of GREML estimate using simulated phenotypes based on UK Biobank genotypes. Bars are 95% CI based on 100 replicates. The unit for the number of SNPs is thousand. This result was based on 858K SNPs (after QC) and 10,000 individuals that were randomly selected from UK Biobank. SNPs in each bin were randomly drawn from the 858K SNPs independently. The number of causal SNPs was 10,000 that were randomly selected in each bin. The true simulated value for the genetic correlation was 0.6 and that for the heritability was 0.5 for both traits. Overlap (0%, 10% and 20%) stands for the percentage of overlapping individuals in the first and second traits.

To explore the stability of the accuracy for both methods, we used two additional genotype data sets without imputation, Wellcome trust case control consortium 2 (WTCCC2^18-21^) and genetic epidemiology research on adult health and aging cohort (GERA^22; 23^), which are publicly available (see Supplemental Methods for detailed data descriptions). We also used UKBB raw (non-imputed) genotype data (UKBBr). We calculated the correlation between the LD scores for the HapMap3 SNPs estimated based on the 1KG CEU reference sample (downloaded from https://data.broadinstitute.org/alkesgroup/LDSCORE/) and those based on in-sample genotype data, i.e. UKBB, WTCCC2, GERA and UKBBr data set (Table 1). We found that the WTCCC2, GERA or UKBBr (raw) genotypes were less similar to the 1KG reference genotypes, compared to the UKBB (imputed) genotypes (noting that UKBB samples had been imputed to the combined data of 1KG reference and UK10K data). Table 2 shows that the SE ratio of LDSC estimate to GREML estimate was higher for WTCCC2, GERA or UKBBr than that for UKBB. Figure 2 shows that the accuracy of GREML was consistent across different data sets, whereas that of LDSC was decreased for WTCCC2, GERA or UKBBr, compared to UKBB data set. This was probably due to higher (or lower) correlation between LD scores based on the 1KG reference and the in-sample genotype data sets (Table 1) which might positively or (negatively) affect the accuracy of LDSC estimates. For WTCCC2, GERA and UKBBr data, the SE ratio of LDSC to GREML based on different number of individuals is shown in Figures S6, S7 and S8.

**Figure 2.**
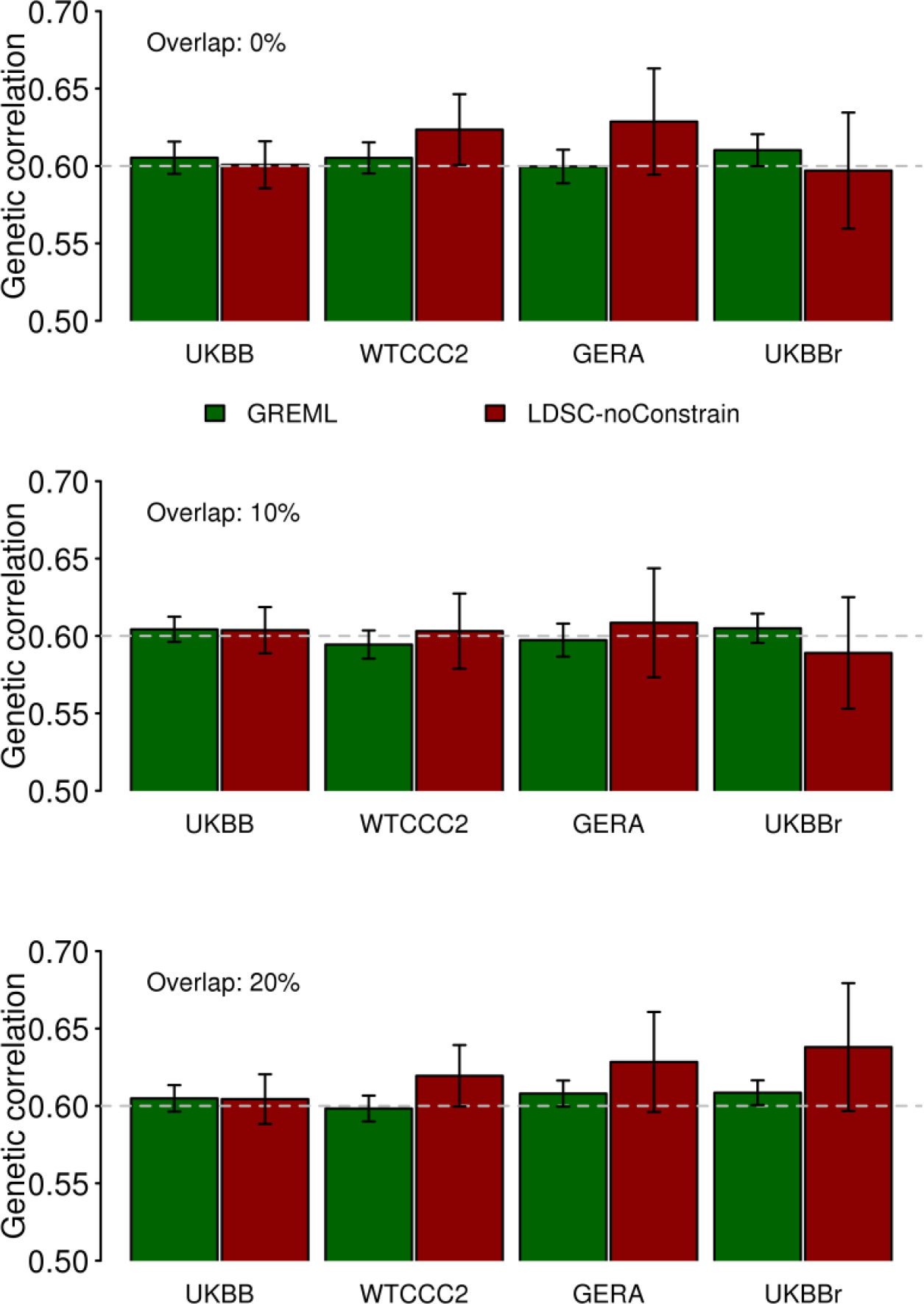
Estimated genetic correlation with GREML and LDSC (without constrain to the intercept) based on different genetic data sets. Simulation was based on 10,000 individuals that were randomly selected from UKBB, WTCCC2, GERA and UKBBr (the raw genotype of UKBB), with 858K, 432K, 239K, and 124K SNPs, respectively. Bars are 95% CI based on 100 replicates. Overlap (0%, 10% and 20%) stands for the percentage of overlapping individuals in the first and second traits. The grey dashed line stands for the true simulated genetic correlation 0.6.

**Table 1.**
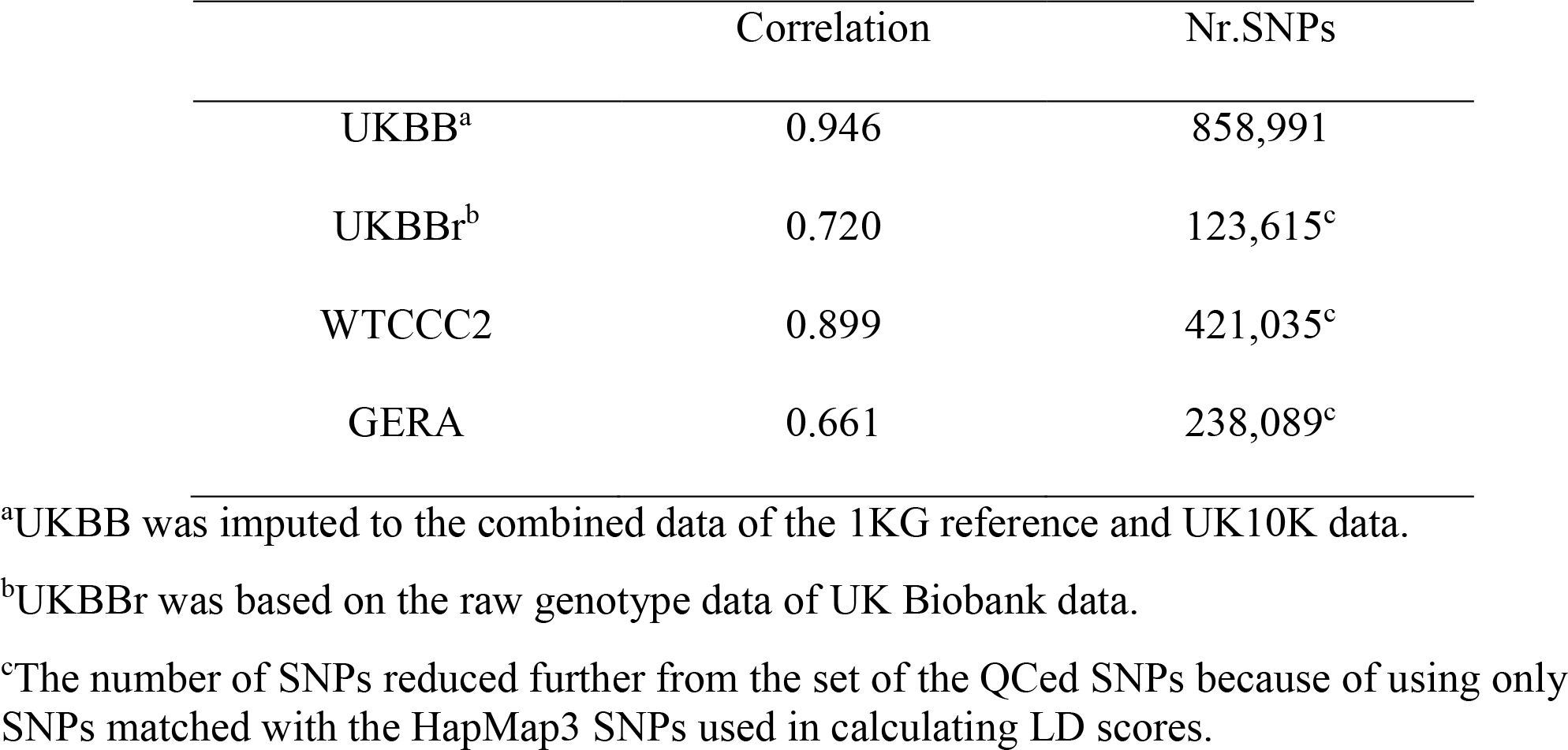
Correlation between LD scores estimated based on the HapMap3 SNPs using the 1KG CEU reference sample and that from different target populations

|  | Correlation | Nr.SNPs |
| --- | --- | --- |
| UKBB<sup>a</sup> | 0.946 | 858,991 |
| UKBB<sup>r</sup><sup>b</sup> | 0.720 | 123,615<sup>c</sup> |
| WTCCC2 | 0.899 | 421,035<sup>c</sup> |
| GERA | 0.661 | 238,089<sup>c</sup> |

<sup>a</sup>UKBB was imputed to the combined data of the 1KG reference and UK10K data.
<sup>b</sup>UKBB<sup>r</sup> was based on the raw genotype data of UK Biobank data.
<sup>c</sup>The number of SNPs reduced further from the set of the QCed SNPs because of using only SNPs matched with the HapMap3 SNPs used in calculating LD scores.

**Table 2.** The ratio of SE of LDSC estimate to that of GREML estimate using simulated phenotypes based on UKBB, WTCCC2, GERA and UKBBr genotypes in the scenarios without overlapping individuals

|  | 800k | 400k | 200k | 100k |
| --- | --- | --- | --- | --- |
| UKBB | 1.60(0.15) | 1.70 (0.18) | 1.85 (0.25) | 2.04 (0.33) |
| WTCCC2 | NA | 2.15 (0.31) | 2.35 (0.43) | 2.68 (0.61) |
| GERA | NA | NA | 2.87 (0.56) | 3.31 (1.17) |
| UKBB<sup>r</sup> | NA | NA | NA | 3.74 (0.79) |

Genome partitioning analyses are an emerging tool to estimate the genetic variance and covariance explained by functional categories (e.g. DNase I hypersensitive sites (DHS) and non-DHS ^24^). Currently, genomic partitioning analyses focus on SNP-heritability enrichment analyses, formally testing for enrichment of signal compared to the expectation that the estimates are proportional to the number of SNPs allocated to each annotation. Considering genomic partitioning in cross-disorder analyses is a natural extension to identify regions where genetic correlations between disorders are highest and lowest. Here, we assessed the performance of the methods in the context of genome partitioning analyses using simulated phenotypes based on UKBB genotype data. A better LDSC approach to estimate genetic correlation for each category might be sLDSC, stratifying by genomic annotation; however, this method is currently under development (i.e. there is software (see Web Resources), but there is no published document or paper verifying the method). Nonetheless, since the sLDSC is available to the research community, we applied both LDSC and sLDSC to estimate partitioned genetic correlations for the simulated data (Supplemental Methods). For genome partitioning analyses, we showed that LDSC estimates of genetic correlation were biased whether using LD-scores estimated from the 1KG reference or in-sample data (UKBB) while GREML estimates gave unbiased estimates for each functional category (Figure 3). sLDSC estimates were unbiased only when using LD-scores from the in-sample data, and their SEs are relatively larger than those of GREML or LDSC (Figure 3). This was probably due to the fact that the different distribution of causal variants and their effects between DHS and non-DHS regions were better captured by an explicit covariance structure fitted in GREML. We also applied the methods to a range of simulation scenarios and found similar results in that GREML performed better than LDSC or sLDSC (Figure S9 and Table S1), which was consistent with the previous results (Figures 1 and 2). It is notable that in a deliberately severe scenario (e.g. causal variants are simulated only within few kb of a boundary) GREML could give biased estimation of genetic correlation ^13; 24^.

**Figure 3.**
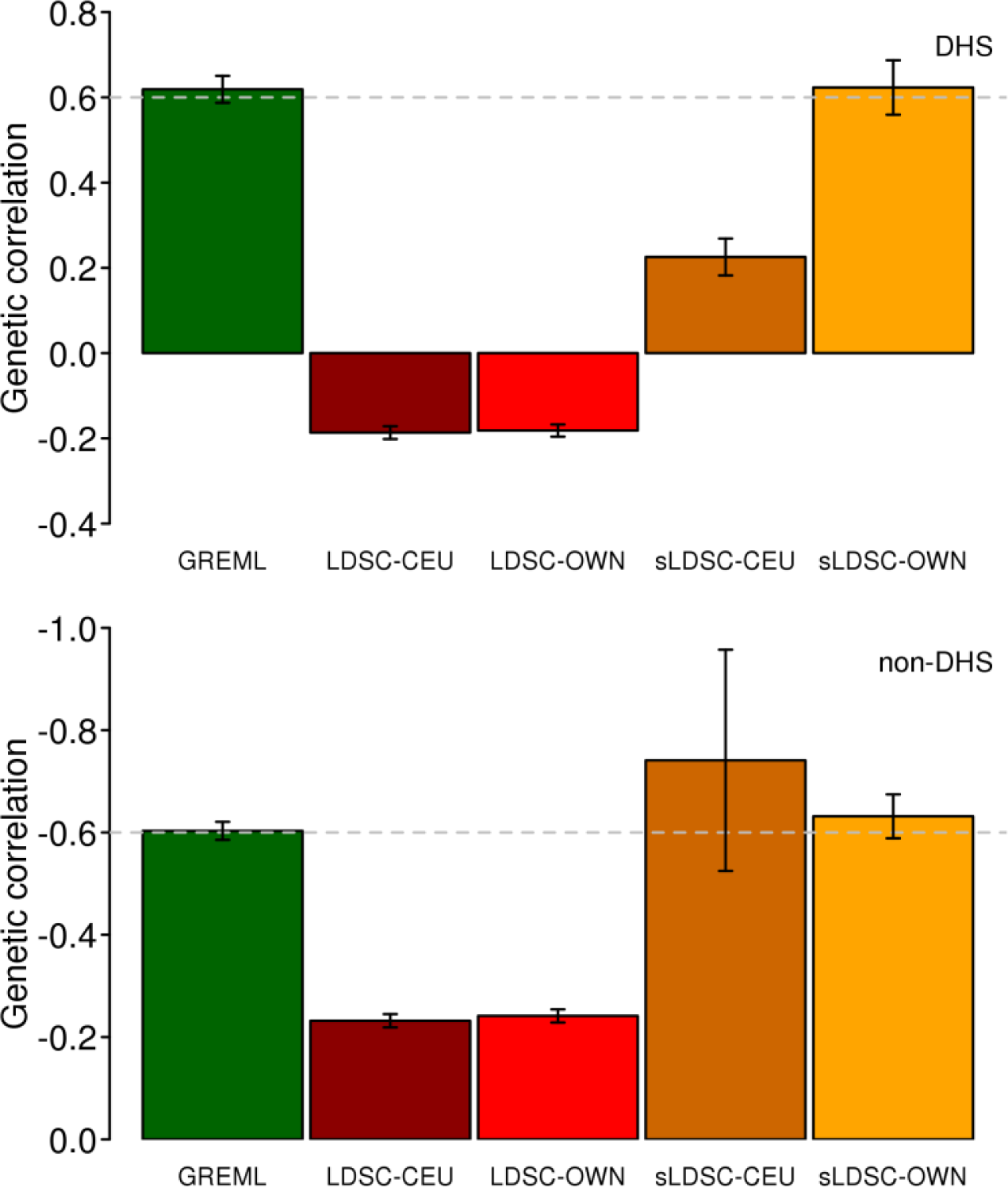
Estimated genetic correlation of simulated data based on a genomic partitioning model. Simulation was based on 10,000 individuals that were randomly selected from UKBB with 858K SNP. Based on Gusev et al.^24^, the 858K SNPs across the genome were stratified as two categories: DHS (194K SNPs with 2268 causal SNPs) and non-DHS (664K SNPs with 7732 causal SNPs). The genetic correlation for the simulated phenotypes between the first and second traits was 0.6 and -0.6 in DHS and non-DHS region, respectively. Bars are 95% CI based on 100 replicates. LDSC-CEU: Using LD-scores estimated from 1KG reference data. LDSC-OWN: Using LD-scores estimated from UKBB. sLDSC-CEU: Using stratified LD-scores estimated from 1KG reference data. sLDSC-OWN: Using stratified LD-scores estimated from UKBB. The presented results were based on 0% overlapping samples between the first and second traits and those based on other scenarios (e.g. 10% and 20%) are presented in Table S1.

While focusing on the accuracy of genetic correlation estimates, there is an important implication for the bias in SNP-heritability estimates for both GREML and LDSC (Figure S10). When using the WTCCC2, GERA and UKBBr data, which were less similar to the 1KG reference genotypes, compared to the UKBB data, LDSC estimates were substantially biased whereas GREML estimates were close to the true value in estimation of SNP heritability (Figure S10). However, this result is well known and LDSC was not recommended for SNP heritability by the original authors ^10^, but rather for relative enrichment analysis. Despite this, LDSC is widely used for SNP-heritability estimation (because it is quick and simple). Thus, for completeness we include analyses for different scenarios to quantify the properties of the methods. When reducing the number of SNPs, estimated SNP-heritabilities from LDSC were consistently unbiased; however, those from GREML were proportionally underestimated (Figure S11). When using non-HapMap3 SNPs, LDSC estimates were consistently biased (Figure S12) and less accurate, compared to GREML estimates (Figures S13 and S14), which probably explains why LDSC is implemented using only HapMap3 SNPs. Although the genetic correlation is robust to such biasedness ^4; 11^, SNP-heritability itself should be carefully interpreted for both GREML and LDSC. We also noted that LDSC and sLDSC estimates for SNP-heritability were biased in the genome partitioning analysis (Figure S15) although the estimated enrichment was close to the true value when using sLDSC and in-sample LD scores (Figure S15).

We used real phenotype and individual genotype data from the Psychiatric Genomics Consortium (PGC) and UKBB to estimate genetic variance and covariance between SCZ and BMI using LDSC and GREML (Table 3 and Figure S16). We also used publicly available GWAS summary statistics for LDSC to see how much the SE of estimates could be reduced by increasing the number of samples and number of SNPs. For real data analyses, we obtained theoretical SE to assess the accuracy of the methods. GREML and LDSC estimates for the SNP-heritability were 0.192 (SE 0.004) and 0.280 (SE 0.016) for SCZ and 0.184 (SE 0.004) and 0.255 (SE 0.014) for BMI. The notable difference between GREML and LDSC was probably because of a relatively small number of SNPs (500K) that might result in underestimated GREML SNP-heritability (see Figure S11). This is one of the caveats of using GREML with real data that usually comprise multiple cohorts genotyped on different platforms, such that, even with imputation, the overlapping set of SNPs imputed with high confidence may be limited. The estimated genetic correlation for GREML and LDSC was -0.136 (SE 0.017) and -0.173 (SE 0.031). This indicated that the GREML estimate was 3.5 and 1.8 times more precise than LDSC estimates for the SNP-heritability and genetic correlation, respectively. For LDSC, we also considered using additional GWAS summary statistics from publicly available resources^25; 26^. The sample sizes used for additional LDSC analyses (LDSC-meta) are summarized in Table 3. The estimated SNP-heritability was 0.259 (SE 0.019) for SCZ and 0.121 (SE 0.007) for BMI, and the estimated genetic correlation was -0.087 (SE 0.019). Although sample size was increased 2.7-fold, the SE of LDSC estimate was not smaller than that for GREML estimate (SE = 0.017 vs. 0.019, and p-value = 4.54E-15 vs. 4.91E-06 for GREML vs. LDSC) (Table 3). It should be noted that GREML estimates used a homogeneous population (within UK and after stringent QC excluding population outliers) whereas LDSC-meta1 and -meta2 were based on combined meta-data sets consisting of ~ 80 different studies for which there is much more uncertainty about homogeneity than when using a single study cohort such as UKBB. The large difference of the estimates between LDSC and LDSC-meta1 (or -meta2) was probably due to the fact that heterogeneity among the 80 different studies resulted in underestimation of the common genetic variance and covariance, and that the difference of LD scores between the target and 1KG reference data would bias the LDSC estimates as shown in Figure S10. We also analysed height data^27^ and found a similar pattern in that GREML estimates were more accurate than LDSC estimates whether using the same data or using additional GWAS summary statistics for LDSC (Figure S17 and Table S2).

**Table 3.**
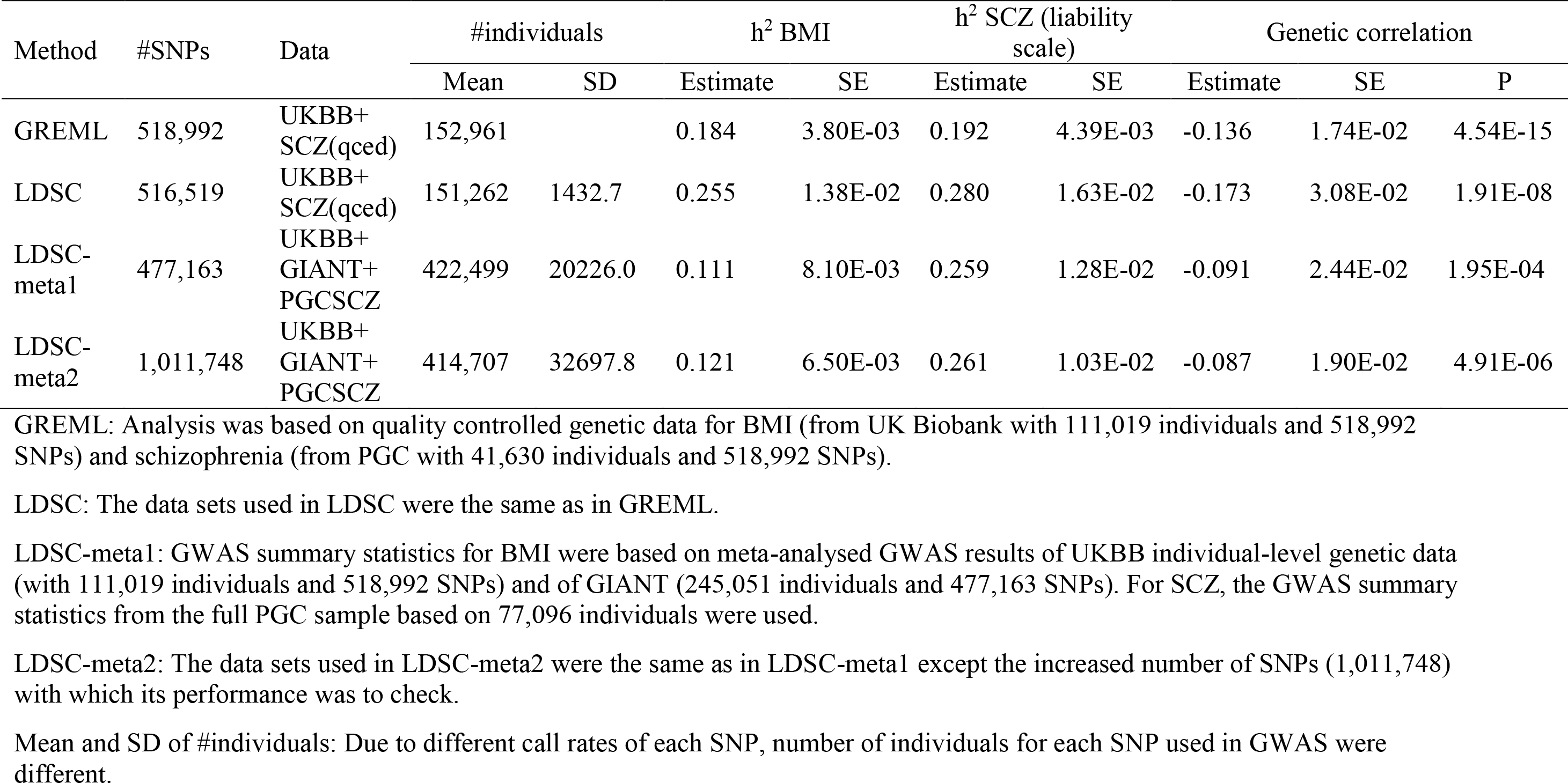
Heritability and genetic correlation based on different data sets

In the real data analyses, we carried out a functional category analysis partitioning the genome into regulatory, DHS, intronic and intergenic regions using GREML (Figure 4 for SCZ/height and Figure S18 for SCZ/BMI). For SCZ and height, the genetic correlation for the regulatory region was negative and significantly different from 0 (p-value = 0.0028; Figure 4). We also compared the results with the LDSC genetic correlation estimation (Figure S19 and S20), and show that the estimates were similar between LDSC and GREML. However, GREML had a lower p-value (0.0028 in Figure 4) than LDSC using LD-scores from the 1KG reference data (p-value = 0.04) or using LD-scores from the in-sample data (p-value = 0.007). We note that current sLDSC software does not provide a SE of estimated partitioned genetic correlation for each category; therefore we did not attempt using the software for the real data analysis. For SNP-heritability estimation, the SE of the estimate for each category was much lower for GREML than sLDSC, ranging from 2.2 to 5.9-fold (Table S3).

**Figure 4.**
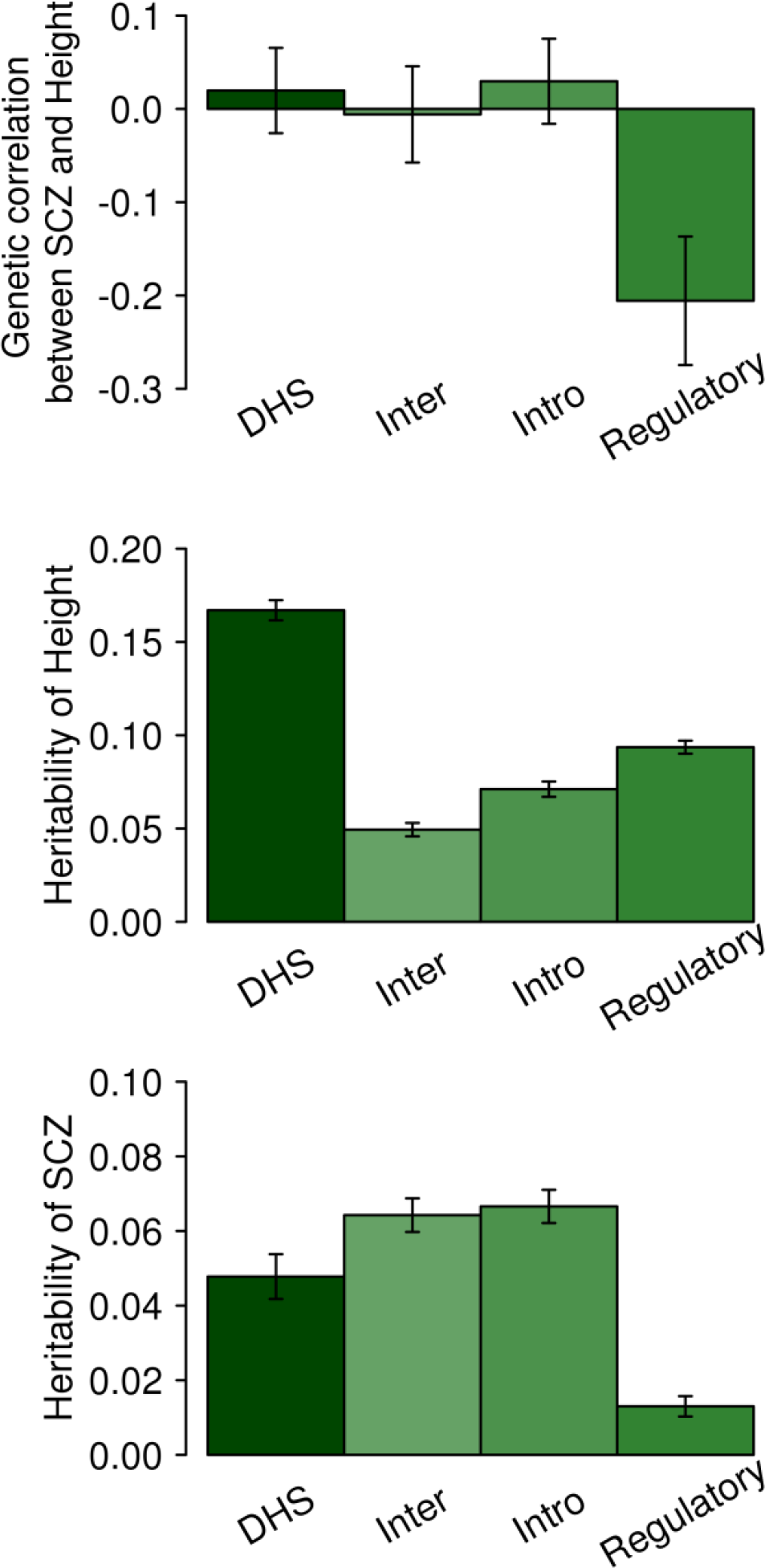
Genetic correlation between SCZ and height and heritability based on SNPs in partitioned genomic regions estimated with GREML. A joint model was applied by fitting four genomic relationship matrices simultaneously, each estimated based on the set of SNPs belong to each of the functional categories (regulatory, intron, intergene and DHS). The bars are standard errors. P-value for the estimate significantly different from 0 was 0.0028, 0.52, 0.91 and 0.67 for regulatory, intronic, intergenic and DHS region, respectively.

### Box 1.

Summary points

1. GREML and LDSC can both provide unbiased estimates of the genetic correlation between two traits. GREML requires individual level genotype data, while LDSC requires only association summary statistics and LD scores per SNP. If LD scores have been calculated from the same sample as the association statistics, then GREML and LDSC provide similar estimates of the genetic correlation. However, in practice LD scores are estimated from external reference samples of the same broad ethnicity, which can lead to bias in the estimates (Figure S21 and S22). As a rule of thumb, when LDSC and GREML estimates are dissimilar, we recommend reporting the estimate with a lower SE. The theoretical SE of the estimates is a reliable indicator to determine the better estimator, which agrees well with the empirical SE (from simulation replicates) (Figure S23).
2. When combining multiple data sets to estimate genetic correlations between multiple traits, it is possible, in practice, that the number of SNPs remaining after QC is relatively small. When the number of available SNPs is small, the SE of LDSC estimates for genetic correlation can be increased relatively more, compared to that of GREML estimates (Figure S2).
3. SNP-heritability has a different property, compared to genetic correlation since the latter is robust to biased estimation of genetic variance and covariance (presumably the biases occur in the numerator and denominator and hence approximately cancel out)^4; 11^. Especially when using a small number of SNPs (< 500K) for GREML or when using multiple meta-data sets for LDSC, estimated SNP-heritability itself should be reported with caution as both methods can give biased estimates.
4. When using a study cohort, it is desirable to measure heterogeneity between the cohort and 1KG reference data (e.g. measuring the correlation between LD scores estimated based on the cohort and 1KG reference data as in Table 1). If the correlation is not close to one, LDSC estimates should be carefully interpreted. We recommend that when GWAS summary statistics are provided, cohort specific LD scores are provided also. It is also warranted that an optimal approach to meta-analyse LD scores across multiple cohorts should be developed to improve LDSC performance ^28^.
5. When using extensive meta-data that possibly include heterogeneous sources, there are two problems. Firstly, the LD scores estimated from reference samples such 1KG reference may be a poor representation of the LD scores of the heterogeneous meta-data, such that the accuracy of LDSC decreases. Second, the distribution of causal variants and pleiotropic effects may be different between heterogeneous sources such that the estimates can be biased (capturing only common effects between heterogeneous sources). This implies that LDSC estimates should be reported with caution when using extensive meta-data sets (Table 3).
6. 6. One of advantages of having access to individual-level genotype data comes when more detailed analyses are required, such as genomic partitioning analyses. As shown in Figure 4, a GREML genomic partitioning analysis reveals a significant negative genetic correlation between SCZ and height for the regulatory region, which genome-wide GREML or LDSC approach has less power to detect.

LDSC and GREML are the methods that have been widely used in estimating genetic correlation, shedding light on the shared genetic architecture of complex traits, based on genome-wide SNPs. Two critical parameters for assessing methods are bias (whether the estimates over replicated analyses differ from the true value) and accuracy (reflected by the standard error of the estimate). Although the property of the accuracy of GREML has been thoroughly studied and tested ^29; 30^, that of LDSC has not been sufficiently investigated. In this report, we compare the accuracy of GREML and LDSC estimates based on various scenarios using simulated as well as real data sets, and draw simple but useful guidelines (Box 1).

Both GREML and LDSC are methods that aim to estimate the same genetic correlation parameter based on genetic variants across the allelic spectrum as defined earlier and the definition is invariant across the methods. The estimates from both GREML and LDSC are valid if all required assumptions are met. GREML estimates variance/covariance components based on genetic covariance structure estimated from available (in-sample) individual genotypes; whereas LDSC estimates variance/covariance components based on association test statistics corrected for LD structure inferred from the markers in the reference panel (e.g. 1KG of the same ethnicity). The underlying assumption is that the samples generating the GWAS summary statistics are drawn from the same population as the samples generating the LDSC statistics, but here we showed that there can be LD-structure (LD-scores) differences between in-sample and reference data, which impacts parameter estimations (Tables 1 and 2 and Figure S10).

The reduced computing burden of LDSC over GREML makes it the method of choice for generating a quick overview of the genetic relationship between disorders (Table S4). However, our results suggest that important associations could be overlooked. For example, Bulik-Sullivan et al.^11^ reported a negative genetic correlation between BMI and SCZ estimated by LDSC (Estimate = -0.095, SE = 0.025 with p-value = 1.75E-4) which was not significant after Bonferroni correction for the multiple testing. Because of the limited power from LDSC analysis, the shared genetic architecture between BMI and SCZ, perhaps, has had less attention than it is due. We confirmed the negative genetic correlation between BMI and SCZ with a greater confidence (Estimate = -0.136, p-value = 4.54E-15) using GREML. A second example is in analyses investigating the shared genetic architecture between height and SCZ, in which epidemiological evidence points to a negative association ^31^, supported by genetic analyses ^32^. However, there was no evidence of genetic correlation between height and SCZ in whole-genome level analyses of Bulik-Sullivan et al. ^11^ (Estimate = -0.002, SE = 0.022). We used a GREML genomic partitioning analysis and found a significant negative genetic correlation between height and SCZ for the regulatory region (Figure 4). It was noted that the regulatory region was highly enriched for height (Estimate = 0.094, p-value = 7.60E-92 in Table S3), which intuitively supports a significant genetic correlation with SCZ for the region. As shown in Figure 3 and Figure S15, the GREML estimate was closer to the true values with a lower SE than LDSC or sLDSC estimate in simulated data. For the real data analyses (Table S3), GREML had more accurate SNP-heritability estimates (lower SE) than sLDSC. Moreover, the sum of each category matched well with the estimate of the whole-genome for GREML whereas this was not the case for sLDSC (Tables S3).

Here we focused on genetic correlation estimates, and did not consider a number of alternative approaches that have been explored in detail for estimation of SNP-heritability, e.g. LDAK approach^33^, Weighted genomic relationship matrix^34^, MAF stratified^29^ and LD-MAF stratified approaches ^35^. It was beyond the scope of our study to assess if biasedness and accuracy can be improved with these methods, although a general observation is that biases in SNP-heritability estimation can ‘cancel’ in estimates of genetic correlations, as biases impact both the numerator and denominator of the genetic correlation quotient^4; 11^. We note that while under review, two new methods to estimate stratified genetic correlations via GWAS summary statistics ^36; 37^ have been published as alternatives to sLDSC. Those approaches also need external reference samples to infer LD-structure in the actual sample, implying the same problem as for LDSC (#4 and 5 in Box 1). However, to partially address this problem one method ^36^ achieves smaller standard errors than sLDSC through a block diagonalization of the LD matrix. A further study is needed to make explicit comparisons with GREML.

In conclusion, LDSC may be the best tool for a massive analysis of multiple sets of GWAS summary statistics in estimating genetic correlation between complex traits, because of its low computing burden and because summary statistics may be available for much larger sample sizes than those with individual genotype data. However, LDSC estimates should be carefully interpreted, considering the summary points (Box 1). Any interesting findings from LDSC analyses should be followed up, where possible, with more detailed analyses using individual genotype data and with GREML methods, even though sample sizes with individual genotype data may be smaller.

## SUPPLEMENTAL DATA DESCRIPTION

The Supplemental Data include 23 figures, four tables, supplementary methods, consortium members and affiliations, and supplementary references.

## CONSORTIA

The members of the Schizophrenia Working Group of the Psychiatric Genomics Consortium are Stephan Ripke, Benjamin M. Neale, Aiden Corvin, James T. R. Walters, Kai-How Farh, Peter A. Holmans, Phil Lee, Brendan Bulik-Sullivan, David A. Collier, Hailiang Huang, Tune H. Pers, Ingrid Agartz, Esben Agerbo, Margot Albus, Madeline Alexander, Farooq Amin, Silviu A. Bacanu, Martin Begemann, Richard A. Belliveau Jr, Judit Bene, Sarah E. Bergen, Elizabeth Bevilacqua, Tim B. Bigdeli, Donald W. Black, Richard Bruggeman, Nancy G. Buccola, Randy L. Buckner, William Byerley, Wiepke Cahn, Guiqing Cai, Dominique Campion, Rita M. Cantor, Vaughan J. Carr, Noa Carrera, Stanley V. Catts, Kimberly D. Chambert, Raymond C. K. Chan, Ronald Y. L. Chen, Eric Y. H. Chen, Wei Cheng, Eric F. C. Cheung, Siow Ann Chong, C. Robert Cloninger, David Cohen, Nadine Cohen, Paul Cormican, Nick Craddock, James J. Crowley, David Curtis, Michael Davidson, Kenneth L. Davis, Franziska Degenhardt, Jurgen Del Favero, Ditte Demontis, Dimitris Dikeos, Timothy Dinan, Srdjan Djurovic, Gary Donohoe, Elodie Drapeau, Jubao Duan, Frank Dudbridge, Naser Durmishi, Peter Eichhammer, Johan Eriksson, Valentina Escott-Price, Laurent Essioux, Ayman H. Fanous, Martilias S. Farrell, Josef Frank, Lude Franke, Robert Freedman, Nelson B. Freimer, Marion Friedl, Joseph I. Friedman, Menachem Fromer, Giulio Genovese, Lyudmila Georgieva, Ina Giegling, Paola Giusti-Rodrı´guez, Stephanie Godard, Jacqueline I. Goldstein, Vera Golimbet, Srihari Gopal, Jacob Gratten, Lieuwe de Haan, Christian Hammer, Marian L. Hamshere, Mark Hansen, Thomas Hansen, Vahram Haroutunian, Annette M. Hartmann, Frans A. Henskens, Stefan Herms, Joel N. Hirschhorn, Per Hoffmann, Andrea Hofman, Mads V. Hollegaard, David M. Hougaard, Masashi Ikeda, Inge Joa, Antonio Juliá, René S. Kahn, Luba Kalaydjieva, Sena Karachanak-Yankova, Juha Karjalainen, David Kavanagh, Matthew C. Keller, James L. Kennedy, Andrey Khrunin, Yunjung Kim, Janis Klovins, James A. Knowles, Bettina Konte, Vaidutis Kucinskas, Zita Ausrele Kucinskiene, Hana Kuzelova-Ptackova, Anna K. Kähler, Claudine Laurent, Jimmy Lee Chee Keong, Sophie E. Legge, Bernard Lerer, Miaoxin Li, Tao Li, Kung-Yee Liang, Jeffrey Lieberman, Svetlana Limborska, Carmel M. Loughland, Jan Lubinski, Jouko Lönnqvist, Milan Macek Jr, Patrik K. E. Magnusson, Brion S. Maher, Wolfgang Maier, Jacques Mallet, Sara Marsal, Manuel Mattheisen, Morten Mattingsda, Robert W. McCarley, Colm McDonald, Andrew M. McIntosh, Sandra Meier, Carin J. Meijer, Bela Melegh, Ingrid Melle, Raquelle I. Mesholam-Gately, Andres Metspalu, Patricia T. Michie, Lili Milani, Vihra Milanova, Younes Mokrab, Derek W. Morris, Ole Mors, Kieran C. Murphy, Robin M. Murray, Inez Myin-Germeys, Bertram Müller-Myhsok, Mari Nelis, Igor Nenadic, Deborah A. Nertney, Gerald Nestadt, Kristin K. Nicodemus, Liene Nikitina-Zake, Laura Nisenbaum, Annelie Nordin, Eadbhard O’Callaghan, Colm O’Dushlaine, F. Anthony O’Neill, Sang-Yun Oh, Ann Olinc, Line Olsen, Jim Van Os, Psychosis Endophenotypes International Consortium, Christos Pantelis, George N. Papadimitriou, Sergi Papio, Elena Parkhomenko, Michele T. Pato, Tiina Paunio, Milica Pejovic-Milovancevic, Diana O. Perkins, Olli Pietiläinenl, Jonathan Pimm, Andrew J. Pocklington, John Powell, Alkes Price, Ann E. Pulver, Shaun M. Purcell, Digby Quested, Henrik B. Rasmussen, Abraham Reichenberg, Mark A. Reimers, Alexander L. Richards, Joshua L. Roffman, Panos Roussos, Douglas M. Ruderfer, Veikko Salomaa, Alan R. Sanders, Ulrich Schall, Christian R. Schubert, Thomas G. Schulze, Sibylle G. Schwab, Edward M. Scolnick, Rodney J. Scott, Larry J. Seidman, Jianxin Shi, Engilbert Sigurdsson, Teimuraz Silagadze, Jeremy M. Silverman, Kang Sim, Petr Slominsky, Jordan W. Smoller, Hon-Cheong So, Chris C. A. Spencer, Eli A. Stah, Hreinn Stefansson, Stacy Steinberg, Elisabeth Stogmann, Richard E. Straub, Eric Strengman, Jana Strohmaier, T. Scott Stroup, Mythily Subramaniam, Jaana Suvisaari, Dragan M. Svrakic, Jin P. Szatkiewicz, Erik Söderman, Srinivas Thirumalai, Draga Toncheva, Sarah Tosato, Juha Veijola, John Waddington, Dermot Walsh, Dai Wang, Qiang Wang, Bradley T. Webb, Mark Weiser, Dieter B. Wildenauer, Nigel M. Williams, Stephanie Williams, Stephanie H. Witt, Aaron R. Wolen1, Emily H. M. Wong, Brandon K. Wormley, Hualin Simon Xi, Clement C. Zai, Xuebin Zheng, Fritz Zimprich, Kari Stefansson, Peter M. Visscher, Wellcome Trust Case-Control Consortium, Rolf Adolfsson, Ole A. Andreassen, Douglas H. R. Blackwood, Elvira Bramon, Joseph D. Buxbaum, Anders D. Børglum, Sven Cichon, Ariel Darvasi, Enrico Domenici, Hannelore Ehrenreich, Tõnu Esko, Pablo V. Gejman, Michael Gill, Hugh Gurling, Christina M. Hultman, Nakao Iwata, Assen V. Jablensky, Erik G. Jönsson, Kenneth S. Kendler, George Kirov, Jo Knight, ToddLencz, Douglas F. Levinson, Qingqin S. Li, Jianjun Liu, Anil K. Malhotra, Steven A. McCarrol, Andrew McQuillin, Jennifer L. Moran, Preben B. Mortensen, Bryan J. Mowry, Markus M. Nöthen, Roel A. Ophoff, Michael J. Owen, Aarno Palotie, Carlos N. Pato, Tracey L. Petryshen, Danielle Posthuma, Marcella Rietsche, Brien P. Riley, Dan Rujescu, Pak C. Sham, Pamela Sklar, David St Clair, Daniel R. Weinberger, Jens R. Wendland, Thomas Werge, Mark J. Daly, Patrick F. Sullivan & Michael C. O’Donovan

## ACKNOWLEDGEMENTS

This research is supported by the Australian National Health and Medical Research Council (1080157, 1087889), and the Australian Research Council (DP160102126, FT160100229). This research has been conducted using the UK Biobank Resource. UK Biobank (http://www.ukbiobank.ac.uk) Research Ethics Committee (REC) approval number is 11/NW/0382. Our reference number approved by UK Biobank is 14575. GERA data came from a grant, the Resource for Genetic Epidemiology Research in Adult Health and Aging (RC2 AG033067; Schaefer and Risch, PIs) awarded to the Kaiser Permanente Research Program on Genes, Environment, and Health (RPGEH) and the UCSF Institute for Human Genetics. The RPGEH was supported by grants from the Robert Wood Johnson Foundation, the Wayne and Gladys Valley Foundation, the Ellison Medical Foundation, Kaiser Permanente Northern California, and the Kaiser Permanente National and Northern California Community Benefit Programs. The RPGEH and the Resource for Genetic Epidemiology Research in Adult Health and Aging are described in the following publication, Schaefer C., et al., The Kaiser Permanente Research Program on Genes, Environment and Health: Development of a Research Resource in a Multi-Ethnic Health Plan with Electronic Medical Records, In preparation, 2013. This study makes use of data generated by the Wellcome Trust Case-Control Consortium. A full list of the investigators who contributed to the generation of the WTCCC data is available from www.wtccc.org.uk. Funding for the WTCCC project was provided by the Wellcome Trust under award 076113, 085475 and 090355.

## WEB RESOURCES

LDSC: https://github.com/bulik/ldsc

MTG2: https://sites.google.com/site/honglee0707/mtg2

PGC GWAS results: http://www.med.unc.edu/pgc

GIANT GWAS results: https://portals.broadinstitute.org/collaboration/giant/index.php/GIANT_consortium_data_files UK Biobank: http://www.ukbiobank.ac.uk

WTCCC2: https://www.wtccc.org.uk/ccc2/

GERA: https://www.ncbi.nlm.nih.gov/projects/gap/cgibin/study.cgi?study_id=phs000674.v2.p2s

